# A National Survey of 205 Canadian Biomedical Research Core Facilities Reveals Structural Challenges and Opportunities for Strengthening Canada’s Research Ecosystem

**DOI:** 10.64898/2026.06.29.732987

**Authors:** Brooke Ring, Charles CT Hindmarch, Stephen L. Archer

## Abstract

**Background:** Canadian biomedical core facilities (BCFs) provide researchers with access to advanced tools and unique technical expertise, essential for research. However, their role, sustainability, and impact remain poorly understood. We report on the evolution of model and existing state of Canada’s 205 BCFs, examining challenges and benefits.

**Methods:** Cross referencing of national databases by the Canadian Innovation Fund (CFI), and from data collected by the Canadian Network of Scientific Platforms (CNSP) allowed he identification of BCFs. Hand curation of these lists validated that cores are operational. To ensure cores not listed by CFI/CNSP were captured, research intensive institutions in Canada were independently searched to identify active cores.

**Results:** There are currently 205 active and operational BCFs located across 9 provinces, which can be further stratified into 9 ‘technical domains’ that describe the nature of services they provide. Quebec (80 cores) and Ontario (75 cores) have the highest confluence of BCFs, with Quebec having a higher ratio of cores per capita.

**Conclusions:** While our data establishes the ubiquity of Canadian BCFs, we highlight substantial challenges including sustainability, governance, evaluation and the recognition of support for core scientists. Here, we establish a framework to address these challenges and to inform best practice, to optimize creation of impactful, accessible and functional biomedical core facilities.

## Introduction

Recently, Canada has created definitions to distinguish its research facilities, in an effort to provide clarity for new funding streams^1^. Core facilities are defined as shared centres that provide access to multiple researchers in a local region. Core facilities exist in diverse disciplines including material sciences, physics, chemistry, biochemistry, and computational sciences^2^. Biomedical core facilities (BCFs) are scientific platforms that contain specialized infrastructure and expertise relevant to performing research on the health and diseases of humans and animals. BCFs are accessible to researchers within universities, academic health science centers, industry, government or research centres, typically on a cost recovery basis. BCFs provide individual researchers, known as principal investigators (PIs), and their trainees, access to specialized scientific equipment, services and expertise that complement resources in their own laboratories^3,4^. The type of equipment and expertise available in core facilities are generally too expensive for an individual laboratory to purchase and maintain and/or too specialized to operate^1^. Equipment suited to a BCF includes, but is not limited to: flow cytometry, genomic sequencers, mass spectrometers, advanced microscopes (light, confocal, super resolution, and electron microscopes), high-fidelity imaging platforms for physiological analysis of animal models of human disease (echocardiography, micro-PET-SPECT-CT or MRI scanners), animal care facilities, spatial omics platforms, and histology services. A BCF’s equipment is operated and maintained by career scientists called *core scientists* or *platform scientists*, who typically possess advanced degrees, but do not have individual research laboratories. Core facility organizational models vary, ranging from those focused-on training highly qualified personnel (HQP), to others focused on advancing research excellence or prioritizing protocol development and/or revenue generation^5,6^. Depending on their model, instrumentation and personnel within BCFs are usually shared with a broad research community (internal or external to the Institution)^3,4,7,8^.

Over the past two decades, the complexity of scientific research has grown, necessitating a reassessment of the role of core facilities in the Canadian research ecosystem. In this article we will consider whether extending the model that embraces the use of core facilities, staffed by core scientists, would facilitate optimal use of publicly funded research infrastructure and enhance the ability of PIs to perform world class research. With the goal of identifying knowledge gaps to inform the evolution of core facilities, we will examine the role that BCFs have played in advancing research in Canada over the past two decades, highlighting strengths and weaknesses of the current BCF models for funding, human resource management, operations, and evaluation metrics. Recognition of current gaps will inform policies for the establishment and sustainability of core facilities in Canada and beyond.

## Methods

### Eligibility

For this study, biomedical core facilities are defined as shared centres that provide access to multiple researchers, and which possess infrastructure and expertise^1^ classified in one or more of the nine technical domains: microscopy, medical imaging, flow cytometry, histology, biochemistry/structural biology, cell culturing, drug discovery, genomics and the study of preclinical animal models. Facilities were included only if they were located within Canadian academic or research institutions (Supplementary Table 1). Biobanks, institutional animal care facilities, and facilities operating solely within individual investigators’ laboratories were excluded.

### Data Sources

To create a comprehensive catalogue of operational Canadian core facilities, we interrogated two discrete databases, each of which partially describe BCFs. All searches, screening, and data extraction were performed by a single author. First, we used information gathered by the Canadian Network of Scientific Platforms (CNSP)^9^ between the years of 2018 – 2025, which was provided as a.csv file of all the ∼350 Canadian core facilities the CNSP recognize. Cores that did not meet the study definition of a BCF were excluded. Through the CNSP database we captured the province, home institution, and name of the BCF. Next we validated these data points against the facilities and infrastructure reported between 2013 – 2025 on the Canadian Foundation of Innovation’s (CFI) Navigator^5^ web directory. To do this we used the Advanced search filter, and filtered by Province (‘*Alberta*’, ‘*British Columbia’*, ‘*Manitoba*’, ‘*New Brunswick’*, ‘*Newfoundland and Labrador’*, ‘*Nova Scotia’*, ‘*Ontario*’, ‘*Prince Edward Island’*, ‘*Quebec*’, ‘*Saskatchewan*’); then searched each institution (specifically ‘Universities/Colleges/Hospitals’; and under “Sector” we filtered to only include ‘Life sciences, pharmaceuticals and medical equipment’. As neither the CNSP directory nor the CFI Navigator represents a complete national inventory of BCFs, a targeted institutional search was conducted to identify eligible facilities that were absent from both databases, we also searched each of the top 50 research-intensive^10^ (Supplementary Table 1), Canadian institutions in CFI Navigator, to determine if any BCFs were missed from this directory.

### Institutional Validation

Institutional websites within the targeted search were subsequently reviewed to verify BCF operation, classify technology domains offered, obtain BCF contact information, and identify eligible BCFs not captured in the CNSP or CFI Navigator databases. Searches were performed using the surveyed institutions websites’ search function or Google site search, using the terms “scientific platform”, “core facility”, “core facilities”, and “research services”. Search results for each institution were conducted iteratively until no additional eligible BCFs were identified. BCFs were considered operational if institutional webpages indicated ongoing operations through current service descriptions, staff listings, or user access information. Technology domains were determined based on the type of infrastructure and/or expertise they specifed accessible. The listed manager or main contact for each BCFs was derived from their webbsite and recorded into our database. If a BCF was identified as active and was not recorded in the CNSP or CFI Navigator databases, it was added to our database.

### Data Extraction

For each eligible BCF, the following variables were extracted from the CNSP database, CFI Navigator, and institutional websites: *province, city, institution, facility name, website URL, technology domain(s), primary contact person, and contact email address.* Where information differed between sources, institutional websites were considered the gold standard reference source. Facilities identified through multiple sources were merged into a single record following manual review and validation. The final database represented the integration of all eligible BCFs identified through the CNSP directory, CFI Navigator, and institutional website searches.

## Results

In 2025 we identified 205 active BCFs located across nine of Canada’s ten provinces (Figure 2). These core facilities can be classified based on the technologies and services they offer into 9 technical domains (Figure 3A)^11^. For each core, we determined whether it provided services in one or more of nine technical domains: microscopy, medical imaging, flow cytometry, histology, biochemistry/structural biology, cell culturing, drug discovery, genomics and the study of preclinical animal models (Figure 3B). Excluded from this article were cores that were focused outside of these nine biomedical domains; cores that were no longer operational (as evident by inactive websites, cores whose links and content that had not been updated for more then 3-years); and any core-like resources that were identified as residing within an individual PI’s laboratory. These latter resources are not considered BCFs (even if they contained powerful infrastructure) as they largely operate on a different model, often being accessed through collaboration; rather than cost recovery. The majority of BCFs in Canada offer users a single technology (n=162), most commonly microscopy (n=30). The remaining 43 core facilities offer 2 or more technologies. Only 7 core facilities in Canada provide broad services (offering services in 5-7 technical domains) (Figure 3B).

**Figure 1.**
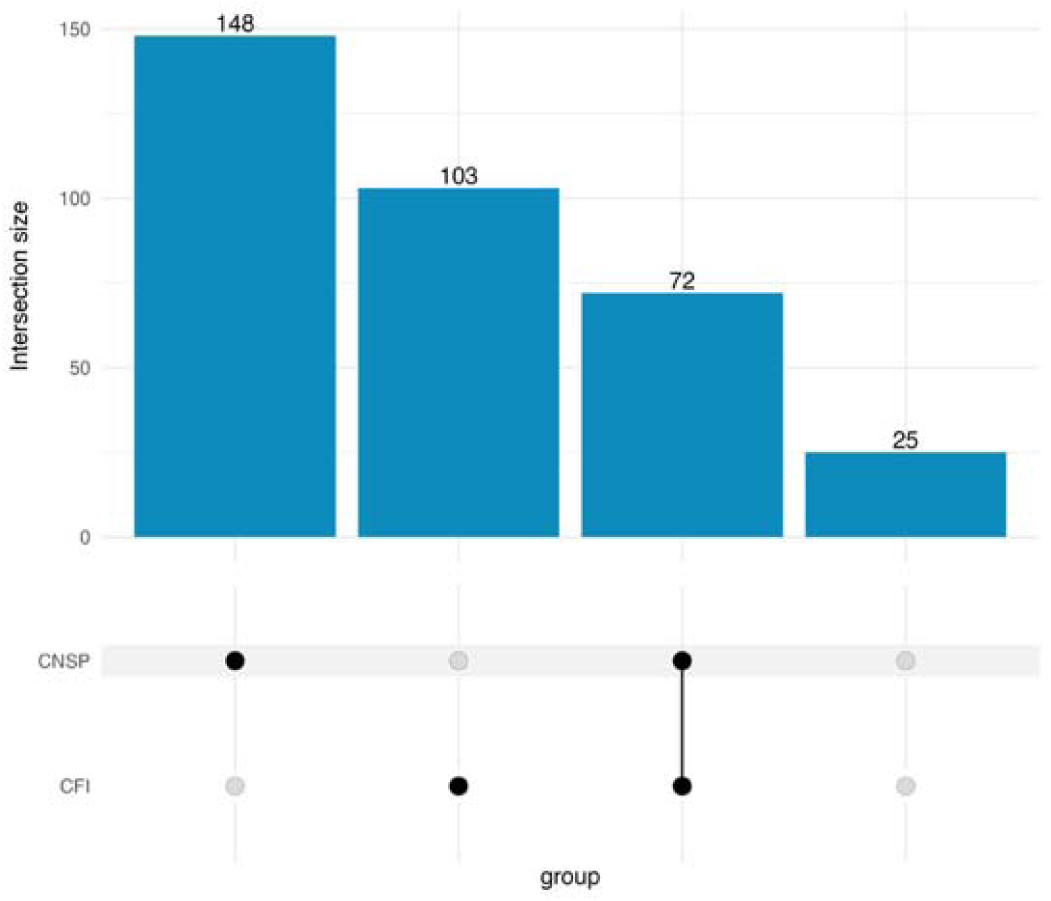
There is no single database for the identification of Canada’s 205 BCFs. Listings of BCF were dispersed across multiple resources. 148 cores were captured by the CNSP database, 103 cores had infrastructure reported on CFI Navigator, with 72 cores recorded on both CNSP and CFI Navigator. The remaining 25 cores were only identified by searching institutional websites.

**Figure 2.**
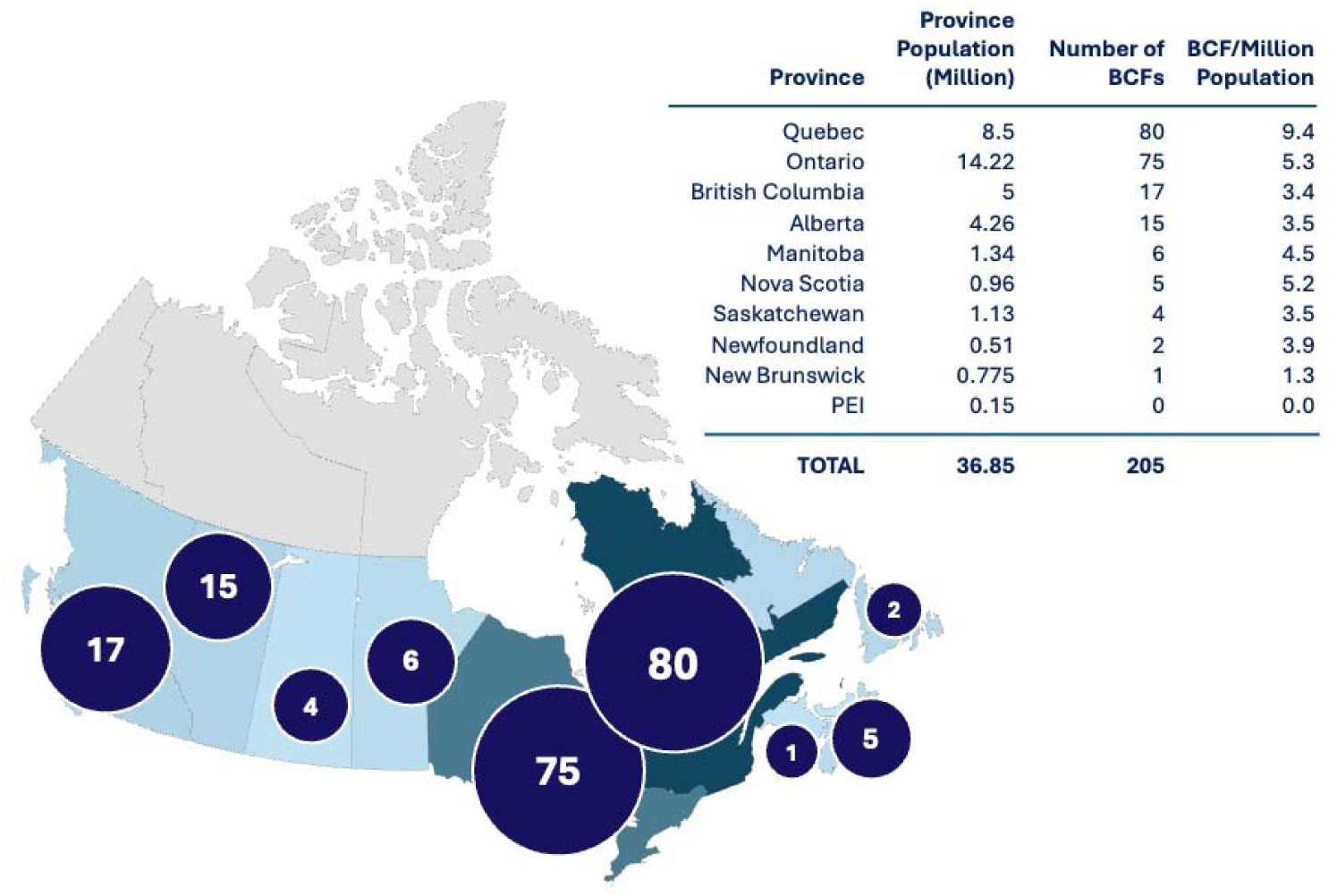
**This maps shows that as of 2025, BCFs (by province) are not distributed uniformly across Canada, when compared to provincial population**^12^. The ratio of BCF/million population provides context for the provincial density of BCFs, displayed per capita. Québec is Canada’s second most populated province, but has the most recognized BCFs (80), likely due to the provinces 2022-2027 Québec Strategy to Support Research and Investment in Innovation (QRIIS2) which is funded by the Québec Infrastructure Plan (QIP) 2026-2036^13^.

**Figure 3.**
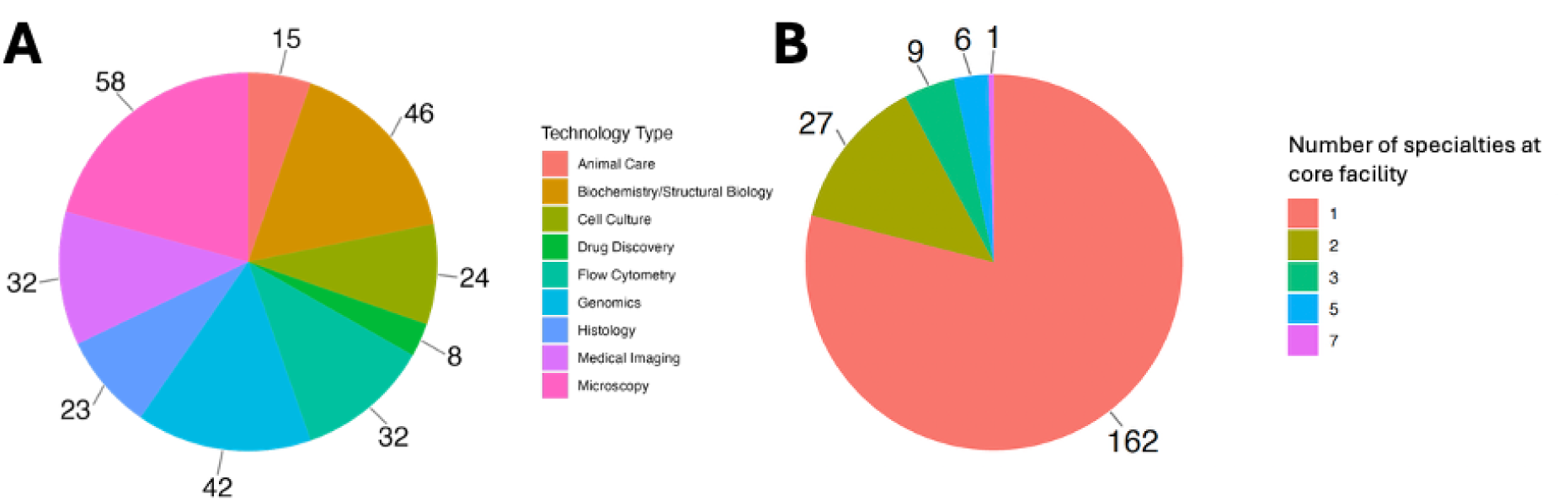
Distribution of technology domains across Canada’s BCFs show that most BCFs offer a single service. **A)** Technical domains of service distribution offered by Canadian BCFs. **B)** Number of technical domains offered by BCFs demonstrates that most BCFs offer services in only 1 technical domain. Data as of April 2025.

## Discussion

Here, we have mined two public databases, and hand curated the results to better understand the scope of the Canadian BCF ecosystem. We will discuss these results within the context of the Canadian scientific funding landscape and how core facilities have evolved over the past two decades, while identifying critical gaps in our understanding of BCF’s impact on Canada’s research landscape.

### Research Funding in Canada

To understand the funding and operation of BCFs it is helpful to review research funding of biomedical science in Canada, which is primarily supported by two key mechanisms: 1) Tri-council funding [made up of three funding agencies the Canadian Institutes of Health Research (CIHR), the Natural Sciences and Engineering Research Council (NSERC), and the Social Sciences and Humanities Research Council (SSHRC)], which covers costs of conducting of research (experimental materials and reagents) and salaries of key scientific staff; and 2) the Canada Foundation for Innovation (CFI), which co-funds acquisition of research equipment and infrastructure platforms at eligible institutions. The CFI typically funds up to 40% of the costs of approved research infrastructure, with the balance being leveraged by funds from provincial funds (40%), and the final 20% provided by industry or non-profit sectors (20%)^14^. Acquisition of a BCF’s research infrastructure is typically funded by CFI grants and the BCF serves to augment research projects of individual PIs, who are funded by Tri-council sources. BCFs are typically utilized by CIHR-funded Individual PIs on a cost recovery (fee for service) basis. While CFI funds were not specifically designated to operate BCFs, most complex, scientific infrastructure is located within CFI-funded core facilities. In 2024, CFI established a new mechanism within their Innovation Fund infrastructure-funding program, called ‘Stream 3’, to fund the creation, renewal and upgrade of *purpose-built* core facilities^14^.

There is currently an undefined and often poorly understood interdependence between funding through Tri-council (like CIHR) and CFI. In reality, there is tremendous interdependence, such that adequate funding in both streams is required for either to be vibrant (Figure 4)^15,16^. While core facilities rely heavily on CFI funding for their creation (acquiring state-of-the-art equipment and infrastructure, as well as construction and renovation costs), cores are equally dependent on PIs, as users, having Tri-council funding (or equivalent) to allow for them to purchase a core’s research services.

**Figure 4.**
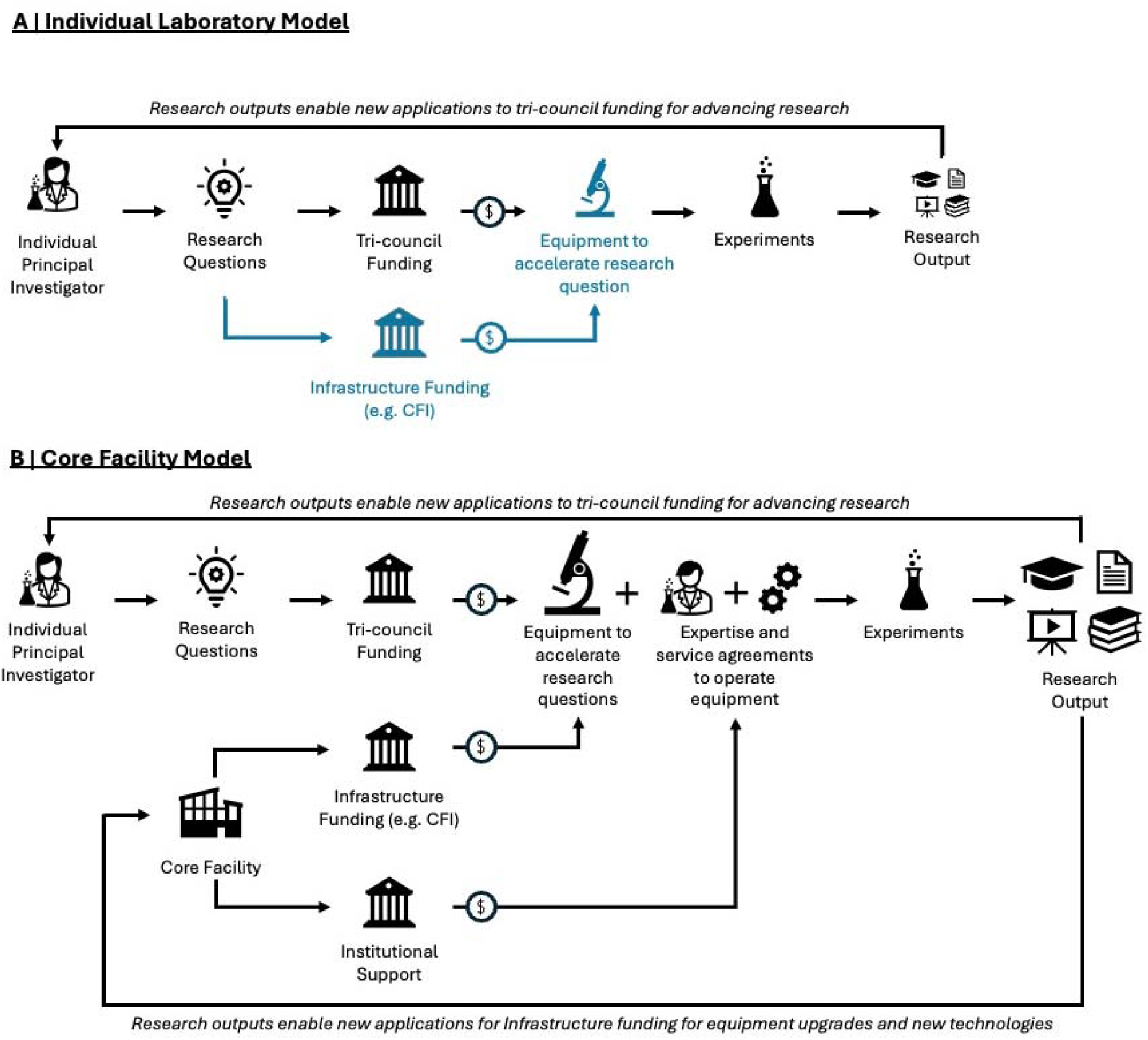
Examples of Canadian research models contrasting the conduct of science by individual PI’s versus individual PI’s supported by access to a core facility. A) The individual laboratory model
shows the traditional route of a PI operating and maintaining equipment in their own laboratory (with optional CFI funding shown in blue). **B) The core facility mode**l demonstrates how BCFs complement the individual laboratory model. With institutional investment and a sustainable funding pathway for scientists (potentially available through a new CFI funding stream), the BCF could constitute a capable and sustainable model.

### Models of Science in Canada

Despite the important interconnection between these two Federal funding agencies (CFI and CIHR), it is not clear what level of coordination there is between the Tri-council and CFI agencies. Understanding the relatively of coordinating operations between these funding agencies is critical if we are to understand the role that core facilities play in Canadian science. Arguably, better coordination between funders of science versus funders of infrastructure may be needed as scientific practice shifts from being the exclusive domain of individual laboratories, working in isolation, to a greater reliance on collaboration amongst PIs and a greater reliance on BCFs to augment a PI’s research capacity^14,17^. Canadian scientists arguably need better access to more complex equipment and with this the specialized scientists that operate these research platforms.

### Individual Laboratory Models

We define the traditional model of science as the *Individual Laboratory Model* (Figure 4A). In this model a PI acquires funding from a Tri-council project grant and uses it to address a specific research question over a period of 5 years^7^. Research output from this funding is a mixture of publications, conference presentations, creation of intellectual property, and knowledge translation, through academic publication and various forms of public engagement. Furthermore, this model supports the education of graduate students and postdoctoral trainees, who conduct research as part of the PI’s research program. If a PI wants to accelerate their research program, they may apply for a CFI grant (e.g., John R. Evans Leaders fund (JELF) or Innovation Fund programs) to purchase state-of-the-art infrastructure that may be used in their own lab or shared with colleagues. The opportunity to apply for these CFI funds is constrained by the individual institution’s CFI funding allocation, which is largely proportional to their Federal funding success^14^. The institution’s total CFI funding envelope defines the maximum amount of funding for which faculty at the institution can apply. A university’s CFI JELF allocation can be used to contribute to PI start-up packages, though at many institutions there is insufficient funding available to meet the needs of all eligible PIs, leaving this source of funding oversubscribed^18,19^. At the time of application for funds, CFI asks for a 5-year budget for the operation and maintenance of the requested equipment (extending from acquisition through the equipment’s lifetime). Because grant budgeting is based on estimates of future use and maintenance cost, there is a risk of creating a deficit when proposed budgets do not cover actual expenses^14^. If awarded, CFI typically provides institutional operating funds (IOF) to support activation and maintenance of the requested equipment^20^, although these funds are often distributed at the discretion of the institution and do not always get passed to the faculty member who secured the CFI funding.

If the limited operational support provided by CFI is insufficient, the service agreements and operating costs must be funded by another mechanism, or the equipment may fall into disrepair. There is no option to reapply for CFI funding for existing pieces of equipment within the *Individual Laboratory Model*^15,20^. Institutions have a fixed envelope of accessible CFI funding and, to be strategic, these funds are typically allocated to new investigators rather than used to sustain or upgrade previously funded infrastructure in a PI’s laboratory^2,21^. This may jeopardize not only the PI’s research program, but the long-term sustainability of investment made by CFI. This issue is important as infrastructure that is properly maintained represents a more effective return on investment for CFI, the institution, the PI and ultimately is better stewardship for the Canadian taxpayer’s investment^4,22^. Unfortunately, the long-term operational status of CFI-funded equipment and core facilities is currently not effectively catalogued and made public and thus it is difficult to quantify objectively. For instance, it is unknown what percentage of CFI-funded facilities remain operational after 5 years.

While *Individual Laboratory Models* have been the predominant traditional structure in academic research in Canada, one could argue that in an era of ‘big science’ where molecular and omics platforms are increasingly complex and expensive to own and operate, cores are needed to supplement the program of individual laboratories^17,23^. In such research areas, the traditional model faces significant challenges, when it comes to the expense of obtaining and maintaining CFI-funded infrastructure. The lack of sustainable funding mechanisms^15,17^, difficulties in attracting and retaining specialized core scientific expertise^18,24^, and inefficiencies in asset inventories cause equipment to be siloed or accidentally duplicated by other laboratories at the same institution, create barriers to scientific progress and reduce the impact of CFI funding. Furthermore, due to the rapid evolution of technology, especially in biomedical science, specialized equipment often rapidly becomes obsolete leading to a need for newer infrastructure. This rapid evolution of scientific platforms and the need to update or replace equipment frequently is hard to sustain in the *Individual Laboratory Model*^25^.

Without financial support beyond the initial grant period, individual researchers face difficult decisions with regards to maintaining operation of research infrastructure. They will often seek institutional or philanthropic funding support^3^, however, many Canadian institutions are often financially constrained and thus struggle to fund cores^24–27^. For example, in Ontario (Canada’s most populous province), tuition was reduced by 10% in 2019 and has remained frozen since, meaning that 2025 budgets have failed to keep up with inflation or negotiated increases in the salaries of staff and faculty. This has created structural deficits in 14 out of 23 universities in Ontario, which limits their ability to support the costs of operation and maintenance of individual PI’s equipment^26–28^.

More commonly, individual laboratories will turn to a model of ‘cost recovery’ to support their equipment. In this model, the PI provides access to their equipment on a fee-for-service basis. This model is different than the traditional collaborative model, in which investigators help each other and share equipment, at no cost, in exchange for shared publication of results. However, the cost recovery model when operating out of a PI’s laboratory quickly confronts the low dollar density of investigators at many major Canadian universities, a term referring to the Federal funding total within a university divided by the number of research-oriented faculty. PIs often rely on a cost-recovery model for their shared equipment, however, without optimal business and accounting practices, and with the limited budgets of most PIs, this is rarely successful^3,4,29^. With limited discretionary funds available to each investigator, PIs are limited in their ability to charge the competitive rates required to cover the true costs of operating high-level equipment. Determining the correct chargeable rate to achieve financial stability solely through cost-recovery must balance both the costs the market will bear and the equipment’s availability (as the PI who acquired the instrument needs to access it to conduct their own research). In addition the rate charged must be competitive to similar equipment within the local environment (especially if there are similar infrastructure facilities running cost-recovery programs)^30^. If the PI charges fees that are too high they run the risk of not getting enough users and thus not recovering a meaningful percentage of their operational and maintenance costs^30^. Another challenge is that service agreements for major equipment increase in price as the equipment ages, typically industry standard is by 7-10%, each year^21,31^. The cost-recovery model, while intended to support research equipment through user fees, is often unsustainable.

Thus, there are challenges to sustaining major research infrastructure within the *Individual Laboratory Model*. While CFI grants provide essential capital investment, the limitations of dedicated funding for ongoing operational and maintenance expenses place a significant burden on individual investigators. Without systematic solutions, researchers often struggle to maintain operation of state-of-the-art technology, ultimately impacting the competitiveness and productivity of their research programs^7^.

### Core Facility Models

#### Organically-formed Core Facility Model

Traditionally cores formed organically, meaning they were born out of infrastructure and services offered within an *Individual Laboratories Models* that provided high quality service while maintaining fiscal stability, through mechanisms including ‘cost-recovery’. These *Organically-formed* core facility models grew by purchasing or inheriting collections of equipment that individual PIs could no longer sustain independently, but which were valuable to the research community^4,32^. To help support their operations these core labs often benefited from departmental funding which supported equipment maintenance and technician salaries. It is common for scientists in such *Organically-formed* core facilities models to comment that they have “accidently found themselves in these roles”^24^. Eventually, the benefit of the equipment and expertise of these cores became embedded into their local scientific community and became vital resources for other researchers^3,7,33,34^. However, in most cases there is no sustainable funding, and the operational, maintenance and personnel costs of these cores constitute a structural deficit within the budget of their home academic department or faculty. Due to the vital importance of *Organically-formed* cores, there is now a growing movement towards establishing *purpose-built* cores, which have the benefits of an *Organically-formed* core but with a proactively sustainable financial operational model. Purpose built cores are the result of vision and strategic planning to create an accessible core with external funding (such as from the CFI), proactively achieve institutional recognition, and implement planning for access and flow that ensures the facility supports the current and anticipated needs of a broad scientific community for a reasonable time frame.

#### Purpose-built Core Facilities

Due to a lack of long-term stability and inability to update or “evergreen” both the *Individual Laboratory* (Figure 4A) and *Organically-formed* core models, there has been a shift toward the establishment of *Purpose-built* core facilities (Figure 4B) within the last decade in Canada^14,17,33^. There are many benefits to establishing a *Purpose-built* core facility, the most compelling being that they are better designed and managed to house state-of-the-art equipment, capable of offering a breadth of services and, based on their boarder scope, more likely to be cost-effective^3^. Funding of research in Canada is on the decline, when adjusted for inflation^18,19,35^ and this has forced PIs to start exploring opportunities to be strategic and cost-effective with their expenditures^15,36^. The establishment *of Purpose-built* core facilities democratizes infrastructure, offering economy of scale, and allows larger numbers of researchers an accessible solution to utilizing specialized equipment and expertise at a subsidized cost, in the sense that they usually do not pay for the equipment or the salary of the core’s staff scientists who operate the equipment^2,3^. In a 2025 article, Jonkman et al, suggested that sophisticated *Purpose-built* core facilities are beginning to mirror the structure of contract research organizations (CROs), which are specialized, fee-for-service research companies that offer research and development services and typically work closely with industry^17^. However, academic, purpose-built cores, unlike CROs, rarely generate a profit and serve to keep costs to researchers low. Additionally, using a *Purpose-built* core allows resources to be made more visible within an institution and thereby reduces inadvertent duplication of equipment, allowing CFI funds to be used to acquire unique pieces of diverse or innovative equipment. If researchers have access to well maintained, professionally operated, innovative technology, their productivity improves^33,37^ (manifesting as increased publication numbers and impact and greater capacity for training HQPs). Moreover, access to such cores increases a researcher’s competitiveness in securing external funding, such as Federal Tri-Council grants, which has the secondary benefit of increasing the CFI envelope for their home institution^4,34,38^. Thus, it can be argued that *Purpose-built* cores not only accelerate research but also expand the research sector both at their home institution and at the national level. Indeed, Universities that fail to invest in BCFs may be left at a disadvantage in the competition for scarce research funds.

Other benefits of purpose-built cores include attraction and retention of the highly trained staff who operate these cores and the PIs who will use these cores to complement their research programs^4,24^. The core scientists are as important to the success of a core as are the infrastructure platforms themselves, since they bring the technology to life and serve as the operators that perform or guide PIs and their labs in using complex equipment^24^. In addition to operating equipment, core scientists often develop and optimize protocols and analyse complex data thus further catalyzing the conduct of science for the individual PIs. Core facilities attract these highly trained staff scientists in part by offering employment benefits that may not exist in the *Individual Laboratory or Organically-formed core* models, including more stable positions, more competitive salaries and benefits, exposure to diverse scientific projects - which makes for a more interesting work experience, and the opportunity to learn new technical and administrative skills^24,25^. *Purpose-built* cores are also better equipped to support the professional development and ongoing training of their scientific staff than are the laboratories of individual PIs. In addition to an improved employment model that attracts and retains HQPs, core scientists in *Purpose-built* cores usually ensure that the users of the core facility receive quality training, and assist users by performing troubleshooting of experimental protocols, developing robust safety practices and ensuring proper data collection, analysis and storage, all of which elevate the quality of science the users receive^24^.

In a 2020 job compensation survey, conducted by the Association of Biomedical Research Facilities (ABRF), core scientists indicated that their job satisfaction comes from a positive work environment, autonomy, and intellectual challenge^34,39^. Well-run core facilities can also be used to attract faculty to an institution by offering services and equipment that constitutes a competitive edge^33,40^, especially attracting early-career researchers (ECR) who are focused on timely generation of data for Federal grant applications. Access to cores allows data generation during a start-up period when these new faculty members are often waiting to occupy and equip their own lab space. Thus, the BCF gives many ECRs a critically important early start. Furthermore, with *Purpose-built* cores expertise taking care of managing and operating research infrastructure, PIs (both junior and senior) are free to focus on experiments that address their research questions, rather than putting effort and time into operating and maintaining complex equipment for which they may have only intermittent need^38^. Core facilities can also benefit researchers by serving as sites for interdisciplinary collaboration, allowing for core scientists to share ideas, connect researchers and be exposed to new techniques to better answer research questions^3,4,38^. For example, when engaging with a BCF, a PI might learn that a core-provided single-cell sequencing, rather than a bulk sequencing platform, might better address their specific research question and thus choose to change their study design to use that equipment. This liberates scientists to use the technology they need; rather than being stuck using the technology they have. Accessible equipment, technical expertise and the facilitation of interdisciplinary collaborations in BCFs inspires and supports new research areas for PIs and offers researchers access to expensive and complex emerging technologies operated by expert scientists^2^.

While the adoption of core facilities in Canada is relatively recent, there are international precedents on which we can draw to inform Canadian best practices for building and operating *Purpose-built* core facilities. Core facilities are mainstays of research productivity in many parts of the world, including France, Germany, the USA and Australia^41–44^.

### International Model for Core Facilities

While many developed nations have established core facility structure and specific career-tracks for core scientists, it is important to note that each country has its own system and the role of and funding for *purpose-built* core facilities varies amongst nations. Australia is a useful comparator to Canada due to similarities in the size of their economies, federal structure, population size and distribution, and heavy reliance on higher education institutions for research output^45^. Thus, this literature review will include a review of Australian biomedical core facilities, to consider their approach to the core facility model^46^.

According to the OECD, Canada and Australia both contribute a GERD of a little less then 2% of GDP. In 2021, Canada invested a GERD of 1.9% of GDP, while Australia invested a GERD of 1.7% of GDP^47^. Despite the lower investment, Australia’s has set a high standard in building world-class core facilities, and according to the QS World University Rankings (Quacquarelli Symonds, a higher education analytics firm), their universities out rank Canadian universities in terms of research productivity and quality, placing nine Australian universities in the top 100, while Canada only has four^48^.

Globally both Canada and Australia R&D outputs are relatively low in comparison to other G8 nations whose GERD investments average 2.7% of GDP^47^; but more specifically, both Canada and Australia have been criticized for their overreliance on the higher education sector as research engines, accounting for more than 35% of the R&D efforts for both countries (Table 1)^45,46^. A common analysis is that overreliance on the higher education sector reflects a structural imbalance, with an under reliance on private business and government sectors, whose contributions are more prominent, productive and create a greater impact in other G8 countries^49^. For example, the United Kingdom in 2021 had a GERD of 2.77% and its investment in higher education was one third that of Canada’s, at only 12% (Table 1).

**TABLE 1.**
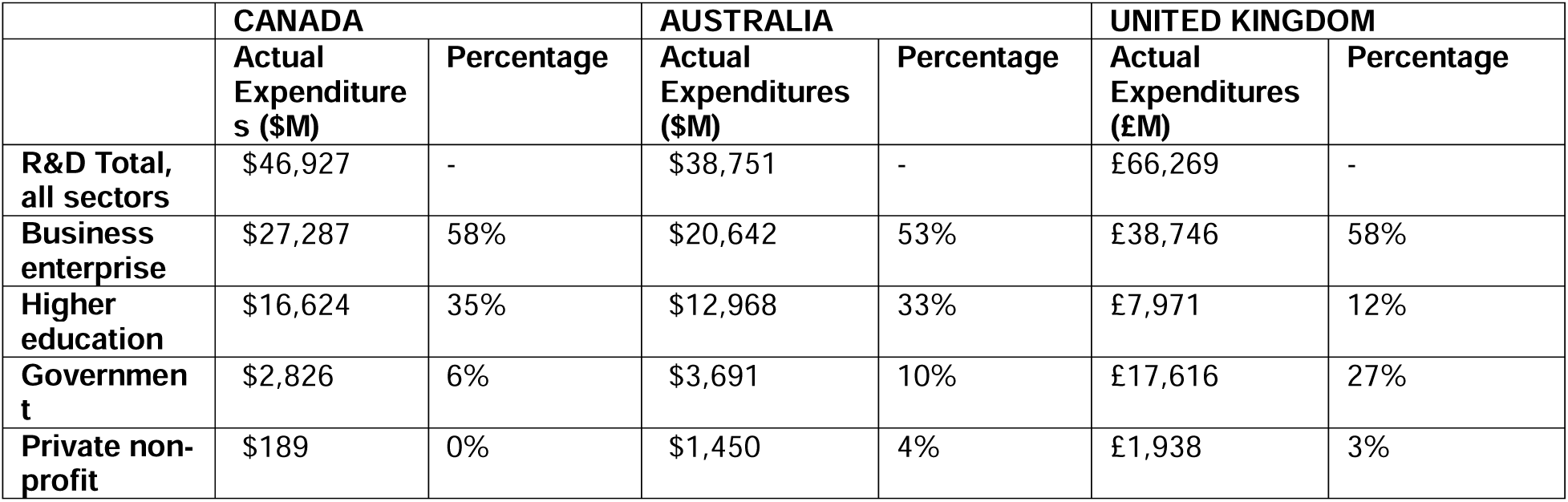
Actual expenditures (millions of dollars (Canadian, Australian and British pounds) of 2021 research and development by sector in Canada, Australia and the United Kingdom. ^50–52^

|  | **CANADA** |  | **AUSTRALIA** |  | **UNITED KINGDOM** |  |
| --- | --- | --- | --- | --- | --- | --- |
| | **Actual Expenditures (\$M)** | **Percentage** | **Actual Expenditures (\$M)** | **Percentage** | **Actual Expenditures (£M)** | **Percentage** |
| **R&D Total, all sectors** | \$46,927 | - | \$38,751 | - | £66,269 | - |
| **Business enterprise** | \$27,287 | 58% | \$20,642 | 53% | £38,746 | 58% |
| **Higher education** | \$16,624 | 35% | \$12,968 | 33% | £7,971 | 12% |
| **Government** | \$2,826 | 6% | \$3,691 | 10% | £17,616 | 27% |
| **Private non-profit** | \$189 | 0% | \$1,450 | 4% | £1,938 | 3% |

Regarding core facilities, the largest difference between the two nations is that Australia has a national strategy for research infrastructure, whilst Canada does not^45^. The Australian Government established the National Research Infrastructure (NRI) program in 2004, which is responsible for Australia’s core facilities, tools, equipment, expertise and other resources that are needed to perform research^53^. The government invested $4 billion over 12 years (from 2018 to 2029) to support the NRI program and make sure it is accessible to Australian researchers and global collaborators. The NRI program is guided by strategic plans called ‘Roadmaps’ statements (which are renewed and published every 5 years)^53^, funded through careful investment plans and enacted through the National Collaborative Research Infrastructure Strategy (NCRIS). Similar to Canada’s CFI program, the NCRIS funds research infrastructure grants and provides services such as a tool to identify critical research networks, like Research Infrastructure Connect^54^, which is similar to CFI Navigator, in that it provides a directory of equipment, data, services and expertise to enable world-leading research to benefit Australians. The NRI’s long-term, strategic approach provides strong foundations for Australia’s research sector, and a national network of facilities that respond to their needs.

Some highlights pertaining to Australian core facilities from the 2021 National Research Infrastructure Roadmap revolved around improving accessibility, funding, workforce development, data management, and industry collaboration. The roadmap suggests a number of potential solutions to explore concerning research infrastructure, such as greater integration, improved funding models, and enhanced digital research infrastructure to support core facilities and maximize their impact^53^.

While Canada currently lacks a national strategy for research infrastructure, there is a pending report on this topic by the Council of Canadian Academies (CCA) which was recently supported by the Strategic Science Fund though the Government of Canada. While, CCA’s reports are not policy decisions, they are designed to inform policy by providing decision makers with the best available evidence^55^.

#### Challenges for Core Facilities in Canada

Despite the benefits of core facilities, there remain significant challenges. Currently, Canada’s core facilities are operating independently (rather than in networks) and are somewhat unregulated, in that there are no established national models, policies, evaluation metrics, or guidelines as to best practices^4,33^. Until recently, Canada lacked a dedicated, national funding mechanism to establish or maintain *Purpose-built* cores, however CFI’s new Stream 3 funding mechanism may remedy these deficiencies. The following section highlights current, key challenges core facilities (including BCFs), are facing.

#### Lack of a standardized, robust financial model

Most core facilities in Canada have reliance on some form of fee-for-service or cost-recovery system, however these revenues rarely cover the facility’s operational and human resources expenses^21^. Without a broad base of regular users (largely Federally funded PIs) there is often not sufficient cost recovery funding to sustain the facility’s resources over time^34^. Some propose that core facilities should consider themselves not-for-profit operators, suggesting that they receive subsidies from their institution or funding agencies, and that most core facilities (depending on their mission) should aim to recover only 30-50% of their budget via cost-recovery^4,29,38^. The target for financial stability must also consider the goal of the core; whether it is to be financially lucrative, break even, or primarily focus on growing science while expecting to incur a deficit that will be covered off by some mix of external and internal base funding. However, there is no consistency or agreement amongst institutions or within government as to the source(s) or magnitude of base subsidies for cores. An important question for Canadian science for the next decade should be whether support for the *Purpose-Built* cores should be primarily from Federal or Provincial sources, the home institution or a mix of all three? When considering funding that might originate from the home institution, should these funds derive from departments, faculties, or from the central institutional budget (the office of the Vice-Principals of Research, VPR)? Some larger Institutions, like the UHN (University Health Network) in Toronto Canada, have restructured their research office to devote staff and resources to support core facilities, using an institutional-level funding budget^56^. Moreover, Canadian universities face an ongoing tension between supporting research excellence, through increasing publications, securing federal funding, attracting and retaining talent, and generating intellectual property versus addressing more basic institutional needs such as infrastructure modernization, maintenance, expansion, and student services^57^. In the absence of objective, standardized metrics to evaluate the scientific and economic impact of core facilities, institutions lack the ability to meaningfully evaluate the return on their investments. This limits their capacity to make strategic, evidence-based decisions about which facilities to support^24,58^. As a result, there is a risk that a Institution’s limited resources are directed toward facilities that are struggling to remain viable, rather than being strategically invested in high-performing cores with the greatest potential to drive research impact and innovation^7,59^.

CFI has been the largest funder of research infrastructure in Canada for almost three decades. Since its inception in 1997 the CFI has funded 14,038 projects with a CFI contribution of $9.7 billion, as of March 2026^60^. As currently structured however, CFI is also not a simple solution to funding core facilities, such as BCFs. Many components must fall into place for successful CFI funding, such as allocating time (∼1.5 years) and resources to prepare a CFI application and gaining internal institution approval to be permitted to apply. Even if an application is put forward by an eligible Institution, the funding success rate is only 39%. We are not suggesting that CFI applications for funding BCF be granted automatically; but rather there should be a peer-reviewed process which evaluates standardized research productivity metrics of cores, assesses best practices and considers a the sustainability of a core’s business model when deciding to renew or replenish core facility funding, upgrade equipment and support BCF personnel.

In the 2023 CFI Innovation competition there were 297 applications submitted, and only 100 were recommended for funding. This represents a 39% success rate, and out of those, only 27% of proposals were in the health research realm (Figure 5A)^61^. Interestingly, in 2023 CFI reported, for the first time, the integration of new infrastructure into core facilities, stating that out of the 100 successful proposals, 51 indicated that some or all the research infrastructure requested would be integrated into a core facility. Considering the estimated CFI investment of $167 million, and an average of 79% of infrastructure actually being integrated into core facilities, approximately 42% of awarded infrastructure would be located in a core facility (Figure 5B)^61^.

**Figure 5.**
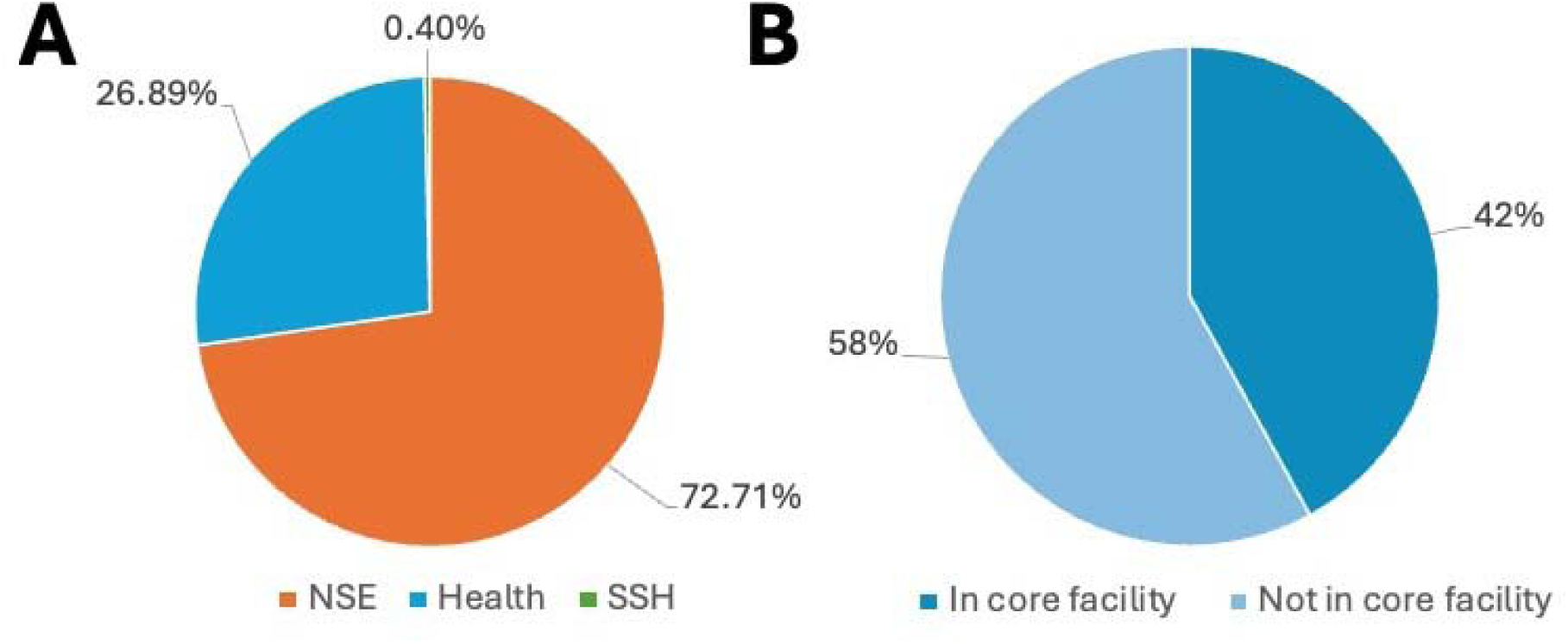
**2023 CFI Innovation fund distribution reported a growing investment of equipment being placed into core facilities**. A) Percentage of total amount awarded by field of research for health, NSE (Natural Science and Engineering) and SSH (Social Sciences and Humanities). B) Percentage of total amount awarded by whether or not the research infrastructure will be integrated into a core facility ^61^

While CFI is a very successful program, it may not be as timely or dynamic as researchers would wish. CFI competitions only occur every 2 years, and if funded, there is a long gap (almost 1.5 years) between application and flow of funds (including provincial matching funds) ^62^. These delays create a potential problem for core facilities and institutions that rely on timely funding both for decisions around procurement and planning of operations. While CFI’s new Stream 3 funding, which began in 2024, could address a number of these challenges and better support establishing, renewing and upgrading core facilities, the decision whether a core director can apply for new or ongoing funding, remains at the discretion of their institution. The role of the institution in determining CFI eligibility is essential, given the need for institutional commitment and matching funding^20^; Thus, incorporating the perspectives of researchers and scientists who lead discovery and innovation in BCFs is important and if properly integrated at the Institutional level could ensure that decisions regarding CFI eligibility are well-aligned with evolving scientific opportunities and the needs of existing BCFs. It also remains unclear whether access to CFI funds will be based on whether the facility is deemed excellent by scientific metrics or whether it will be based on whether the core is a political priority for the institution, who must decide if the core’s alignment to the university/faculty mission merits access to their fixed CFI envelope. Also, provincial matching funds are required for CFI grants, and it remains unclear whether provinces, which currently can provide up to 40% matching funds, will match funding for the new salary support lines within Stream 3 CFI proposals.

#### Poor understanding of impact

Currently there is no agreement on standardization or Key Performance Indicators (KPIs) for the operations of BCFs in Canada^25^,making it difficult to measure the impact or success of a core facility^63^ relative to others in the same institution, province, or nationally. Some core facilities track basic metrics, for example, the number of users, number of trainees, and publications that acknowledge the core’s contributions; however, many core facilities do not even track these rudimentary metrics^37,64^. CFI does require that all awarded Innovation grants submit an annual progress report for the 5 years following the success of their grant. These reports, if replete with accepted metrics, could serve as a rational starting point for more proactive, real-time metric tracking. Currently most metrics are self-reported and are not verified, including number of HQPs trained, research outputs, intellectual property and spin-offs created, national and international collaborations and self-reported benefits to Canadians^20^. The rigour of many of these self-reported metrics, in loosely defined domains such as “benefits to Canadians”, is unclear. Additionally, some core facilities are required to create an annual report for their institutions, however, these annual reports are not standardized in terms of the data reported or the manner of data collection nor are they nationally collected, collated, or published. This lack of standardization and national reporting reflects a missed opportunity for sharing best practices. In a resource constrained environment, it would be helpful to have clear metrics for reporting and measuring scientific and economic impact of BCFs. For example, an independent study by an advocacy group in the United States of America reported by the United for Medical Research (UMR) in 2023 that for every dollar invested in the NIH there was a return of $2.56 in economic activity^65^. Measuring the financial impact of investment by CFI in core facilities would likely reveal that it is a favourable financial investment for the government, but such data are lacking.

Despite these knowledge gaps and in the absence of robust data, CFI is already committed to investing a portion of their 2024-2028 $425 million CAD budget in core facilities through the Stream 3 initiative^14^. Stream 3’s seeks to increase the number of institutions adopting core facilities and implementing formal policies to support them. The 2025 CFI ‘call for proposals’ cites that core facilities have “proven instrumental in attracting, retaining and training top researchers from around the globe”^14^. They also highlighted their potential for fostering collaborations across academic, private, public and not-for-profit sectors. While these concepts, are indeed important for the assessment of the grant proposal, they do not offer detailed benchmarks or KPIs for core facilities^14^.

#### Lack of standard business models for core facilities

As mentioned previously, many core facilities in Canada were founded organically, meaning that they were not purposefully developed with a specific, sound business model at their foundation, but rather evolved with a focus on their scientific purpose rather than long-term stability. Decision makers for most core facilities are often scientists, and while they may possess sound business acumen, they do not typically have formal training in running a business or in budget management^4,7,8^. A case study by Hockberger et al, in 2018, reported on best business practices of NIH-funded core facilities in the USA. They found that institutions found it valuable for their core facilities to create general business models to maximize their services, benefit their users and improve their output (documenting research excellence, training output, and stating profit/loss)^3,29,66^. Core facilities should consider creating multi-year strategic plans and rigorous and transparent budgeting processes. To this end, they might consider adopting popular business models, such as the Business Model Canvas^67^, to understand how to improve, leverage or modify certain aspects of their core facilities through informed decisions. The Business Model Canvas is an introductory business strategy tool that helps map aspects of a business model into nine building blocks: customer segments, value propositions, channels, customer relationships, revenue streams, key resources, key activities, key partnerships, and cost structure^67^. The Business Model Canvas is integral in modern business and informs corporate practices for creating a practical, structured framework to conceptualizing, analyzing and innovating business models^68^. In a special article by Spieth et al from the European Academy of Management (EURAM), it was suggested that the application of this business model had great potential and merit in the field of Research and Development for explaining, developing and operating businesses, like core facilities; though the authors noted that the use of this model for core facilities merits more research^69^. In Canada, the Business Model Canvas is utilized by the MaRS centre (Medical and Research Sciences, North America’s largest Innovation hub) located in Toronto, Ontario, as part of their start-up toolkit for developing a sustainable business model for science-based start-ups^70^.

#### Lack of career track for core scientists

There is currently a disconnect between well-established traditional academic scientists (PIs) ^40,71,72^ and service-oriented (core) scientists^24,28^. Canada has no formalized career track or promotion pathway for core scientists, even though they too may publish, be applicants for awarded extramural grants, and/or teach HQPs^33,40^. In contrast, some jurisdictions outside of Canada have established recognized career tracks for scientist employed by BCFs, including Oregon Health & Science University in the United States^73^, Cambridge University in England^74,75^ or Australia, which is currently in the process of implementing a National level career-track program for research staff and core scientists^53^. There have been 2 internationals surveys by Adami et al in 2021^41^ and Wright et al in 2024 ^24^ with over 500 combined participants which highlighted challenges in creating a career path, gaps in training and the lack of sustainable funding for core scientists.

#### Lack of collaborative networks

Scientific advocacy groups have long played an important role in communicating between government and institutions regarding identifying challenges, evaluating information and disseminating solutions, with the goal of improving research^40,63^. In Canada, these advocacy groups include the CNSP, Canadian Science Policy Centre (CSPC), Genome Canada, BioImaging Canada and the Digital Research Alliance of Canada (DRAC). These organizations represent research interests and support technology development and training^33^. Given financial and governance support, these groups could work together to create national solutions to connect, publicize and support equipment and expertise^24^.

It is unclear where the effective push for dealing with the challenges being faced by BCFs in Canada will come from, grassroots-up or government-down. It is likely that innovation related to BCFs will originate from the most successful cores themselves or perhaps from Canada’s more research-intensive institutions, recognizing the need for efficient policies and best practices to maintain and grow their core facilities. Alternatively, a refresh of what it means to be a core facility and receive funding could be mandated, top down by CFI, as the major funder of this infrastructure in Canada.

#### Identified Knowledge Gaps

This article constitutes a scoping report that identifies current practices and critical knowledge gaps, while considering the repercussion of our currently largely unregulated and unassessed approach to BCFs. The following gaps in the core facilities program in Canada include lack of:

- A catalogued or publicized operational national catalog of core facilities and their equipment.
- Established, standardized, “best practices” guidance for cores in domains including: optimal core model, operational policies, core evaluation metrics/KPIs, core reporting, benchmarking and standardization of cores.
- Sustainable funding for core facilities and their scientists.
- Business training for career scientists who operate or direct cores
- Recognized and funded career tracks for career core scientists.
- Ongoing funding provincial or federal (or institutional) to promote the long-term sustainability of successful biomedical infrastructure cores.
- Clear reporting systems, making the financial impact and return on investment of CFI’s investment in core facilities to the Canadian economy more explicit.
- Support for scientific advocacy networks which enhance the connection between the conduct of science and public policy.
- An explicit, national-level infrastructure strategy for supporting acquisition of scientific equipment and enhancement of scientific technical expertise.
- Governance strategies for cores which would address risk management, operational efficiency, equipment maintenance and upgrading and allow ongoing procurement to deal with rapid technological advances that require equipment updates that cannot be supported by the original core establishment grant^3,4^.

In conclusion, BCFs are central to contemporary research, providing advanced infrastructure, specialized expertise, and collaborative environments that amplify the capabilities of investigator-led laboratories^4,7,76^. This article highlights key challenges across the Canadian landscape, including sustainability, governance, evaluation, and the recognition and support of core facility personnel^3,4,21,22,59^.

Addressing these gaps will require coordinated policy and operational frameworks for best practices, including standardized metrics for evaluation, sustainable funding models, and defined support for core scientists. Despite substantial financial investment, the impact of core facilities remains poorly characterized. Establishing robust approaches to quantify and communicate their scientific and translational contributions will be essential to ensuring proper investment of public funding, long-term sustainability and maximizing the role of BCFs and their staff within Canada’s research ecosystem.

## Acknowledgements

The authors would like to acknowledge Ben Pieter Wood for providing advice on creation and analysis of the database catalogue and providing a figure, and Dr. Doug Quilty and Dr. Jennifer Chen for reviewing and advising on early manuscripts. This study was funded through the Bruce Mitchell Research Program at Queen’s University.

## Authors Contributions

BR and SLA conceived the article topic and structure. BR performed the literature review, conducted the complication of the Canadian Biomedical Core Facility database, analyzed the results, created the figures and drafted the manuscript. CCTH and SLA critically revised the manuscript and advised on content.

## Materials & Correspondence

All inquiries should be directed to Dr. Stephen Archer

