## Supplemental Table 1 for "A National Survey of 205 Canadian Biomedical Research Core Facilities Reveals Structural Challenges and Opportunities for Strengthening Canada’s Research Ecosystem"

**Supplementary Table 1:** Top 50 research intensive Institutions in Canada according to Research InfoSource Inc. ^1^

| **1** | University of Toronto^+^ |
| --- | --- |
| **2** | Université de Montréal |
| **3** | McGill University |
| **4** | University of British Columbia |
| **5** | [University of Calgary](https://researchinfosource.com/r/66e94fbc857571036be3e21f62b1a1e1?eid=2) |
| **6** | [University of Alberta](https://researchinfosource.com/r/efb675d3ead3ef8cbe80a34fdaa4f940?eid=2) |
| **7** | [University of Ottawa](https://researchinfosource.com/r/ab015c1bc40c07cf07823f21a4a3fccb?eid=2) |
| **8** | [Université Laval](https://researchinfosource.com/r/292bc7b74141d02fc7c536c0dce68377?eid=2) |
| **9** | [McMaster University](https://researchinfosource.com/r/34872cdc9c08c622255b049ef95b2f8b?eid=2) |
| **10** | [University of Saskatchewan](https://researchinfosource.com/r/0b4cb3c9c635cf415aede3fbb52ad1ba?eid=2) |
| **11** | Université de Sherbrooke |
| **12** | Western University |
| **13** | University of Manitoba |
| **14** | Dalhousie University |
| **15** | [University of Waterloo](https://researchinfosource.com/r/694faaa4c2e0aba5d69a4954acc9c74f?eid=2) |
| **16** | Queen's University |
| **17** | University of Guelph |
| **18** | University of Victoria |
| **19** | Simon Fraser University |
| **20** | Memorial University of Newfoundland |
| **21** | [York University](https://researchinfosource.com/r/3d8fec37ba5208ae734c8c44e5c76223?eid=2) |
| **22** | Université du Québec à Montréal |
| **23** | [Carleton University](https://researchinfosource.com/r/5642a4cbebced25cee4d65c5178d2fca?eid=2) |
| **24** | Concordia University |
| **25** | [Toronto Metropolitan University](https://researchinfosource.com/r/320051ecf520ab68c5d7115bd55bf18c?eid=2) |
| **26** | Institut national de la recherche scientifique |
| **27** | École de technologie supérieure |
| **28** | University of New Brunswick |
| **29** | Université du Québec à Trois-Rivières |
| **30** | [University of Windsor](https://researchinfosource.com/r/df251730af65872a8eb70b43228fb1df?eid=2) |
| **31** | University of Regina |
| **32** | Université du Québec à Chicoutimi |
| **33** | Université du Québec à Rimouski |
| **34** | [Ontario Tech University](https://researchinfosource.com/r/0810717b6c577298670ece5b8f48317b?eid=2) |
| **35** | [Lakehead University](https://researchinfosource.com/r/72950221c90bb0faa445f74f265b2048?eid=2) |
| **36** | Laurentian University |
| **37** | Université du Québec en Abitibi-Témiscamingue |
| **38** | Wilfrid Laurier University |
| **39** | University of Northern British Columbia |
| **40** | University of Lethbridge |
| **41** | Royal Military College of Canada^++^ |
| **42** | Brock University |
| **43** | [Université de Moncton](https://researchinfosource.com/r/9ce6d867f3b7bab31d64c90b503ebf1f?eid=2) |
| **44** | Université du Québec en Outaouais |
| **45** | Trent University |
| **46** | Saint Mary's University |
| **47** | University of Winnipeg |
| **48** | University of Prince Edward Island |
| **49** | St. Francis Xavier University |
| **50** | Université TÉLUQ |

1.	Canada’s Innovation Leaders. In: Research Infosource Inc.; 2025.
